# Timing gone awry: distinct tumour suppressive and oncogenic roles of the circadian clock and crosstalk with hypoxia signalling in diverse malignancies

**DOI:** 10.1101/556878

**Authors:** Wai Hoong Chang, Alvina G. Lai

## Abstract

The circadian clock governs a large variety of fundamentally important physiological processes in all three domains of life. Consequently, asynchrony in timekeeping mechanisms could give rise to cellular dysfunction underpinning many disease pathologies including human neoplasms. Yet, detailed pancancer evidence supporting this notion has been limited. In an integrated approach uniting genetic, transcriptomic and clinical data of 21 cancer types (n=18,484), we interrogated copy number and transcript profiles of 32 circadian clock genes to identify putative loss-of-function (Clock^Loss^) and gain-of-function (Clock^Gain^) players. Kaplan-Meier, Cox regression and receiver operating characteristic analyses were employed to evaluate the prognostic significance of both gene sets. Clock^Loss^ and Clock^Gain^ were associated with tumour-suppressing and tumour-promoting roles respectively. Downregulation of Clock^Loss^ genes resulted in significant higher mortality rates in five cancer cohorts (n=2,914): bladder (P=0.027), glioma (P<0.0001), pan-kidney (P=0.011), clear cell renal cell (P<0.0001) and stomach (P=0.0007). In contrast, patients with high expression of oncogenic Clock^Gain^ genes had poorer survival outcomes (n=2,784): glioma (P<0.0001), pan-kidney (P=0.0034), clear cell renal cell (P=0.014), lung (P=0.046) and pancreas (P=0.0059). Both gene sets were independent of other clinicopathological features to permit further delineation of tumours within the same stage. Circadian reprogramming of tumour genomes resulted in activation of numerous oncogenic pathways including those associated with cancer stem cells, suggesting that the circadian clock may influence self-renewal mechanisms. Within the hypoxic tumour microenvironment, circadian dysregulation is exacerbated by tumour hypoxia in glioma, renal, lung and pancreatic cancers, resulting in additional death risks. Tumour suppressive Clock^Loss^ genes were negatively correlated with hypoxia inducible factor-1A targets in glioma patients, providing a novel framework for investigating the hypoxia-clock signalling axis. Loss of timekeeping fidelity promotes tumour progression and influences clinical outcomes. Clock^Loss^ and Clock^Gain^ may offer novel druggable targets for improving patient prognosis. Both gene sets can be used for patient stratification in adjuvant chronotherapy treatment. Emerging interactions between the circadian clock and hypoxia may be harnessed to achieve therapeutic advantage using hypoxia-modifying compounds in combination with first-line treatments.

## Introduction

Circadian timekeeping is an essential biological process that influences most, if not all, aspects of eukaryotic and prokaryotic physiology. When its fidelity is compromised, circadian dysregulation may not only result in increased disease risks but could also have an effect on patients’ response to therapy. The core oscillator is coordinated by a set of interlocking transcriptional-translational feedback loop. CLOCK and BMAL1 heterodimerise and bind to E-box elements in PERs and CRYs to drive their rhythmic transcription(1–3). PER and CRY proteins, in turn, inhibit the CLOCK-BMAL1 complex and inhibition is released upon PER and CRY proteolytic degradation(4–6). Along with additional epigenetic and post-translational modification processes, core clock proteins within the suprachiasmatic nucleus serve to sustain the oscillations of peripheral clocks and rhythmic expression of downstream targets(7).

The effect of circadian asynchrony in tumorigenesis was first reported in the 1960s(8). Studies in the 1980s demonstrated that endocrine rhythm disruption could accelerate mammary tumour growth in rats(9, 10). Numerous studies have since shed light on the role of the circadian clock in cancer development in humans. Shift working is thought to be a carcinogen because of dysregulated circadian homeostasis(11, 12). Circadian dysfunction is also thought to be a cancer risk factor in many organ systems(13–19). Aberration in circadian homeostasis is also linked to poor performance in antitumour regimes(17, 20, 21).

Circadian dysregulation is widespread in cancer, yet, tumour-specific abnormalities of clock genes are far from being understood. To help unravel the intricacies of circadian regulation in diverse cancer types, we employed a comparative approach triangulating genetic, transcriptomic and clinical data to discover molecular underpinnings of circadian dysregulation and their effects on patient prognosis. Our pan-cancer integrated analyses demonstrated that the circadian clock had both tumour-promoting and tumour-suppressing qualities that were cell-type dependent. Tumour hypoxia is linked to disease aggression and therapeutic resistance(22). Hypoxia inducible factors (HIFs), the master regulators of hypoxia signalling, are transcription factors containing PER-ARNT-SIM domains and are structurally analogous to core clock proteins BMAL1 and CLOCK(23–25), suggesting that both pathways can be co-regulated. Indeed, past reports have shown that hypoxic responses are gated by the circadian clock(26, 27). We found that the crosstalk between tumour hypoxia and circadian dysregulation harboured clinically relevant prognostic information. Overall, we demonstrated that circadian reprogramming of tumour genomes influences disease progression and patient outcomes. This work could provide a key staging point for exploring personalised cancer chronotherapy and potential adjuvant treatment with circadian- and hypoxia-modifying drugs to improve clinical outcomes.

## Methods

We retrieved 32 circadian clock genes, which included core clock proteins from the Kyoto Encyclopaedia of Genes and Genomes (KEGG) database listed in Additional file 1.

### Cancer cohorts

Datasets generated by The Cancer Genome Atlas were downloaded from Broad Institute GDAC Firehose(28). Genomic, transcriptomic and clinical profiles of 21 cancer types and their nontumour counterparts were downloaded (Additional file 2).

### Copy number alterations analyses

Firehose Level 4 copy number variation datasets were downloaded. GISTIC gene-level tables provided discrete amplification and deletion indicators(29). Samples with ‘deep amplifications’ were identified as those with values higher than the maximum copy-ratio for each chromosome arm (> +2). Samples with ‘deep deletions’ were identified as those with values lower than the minimum copy-ratio for each chromosome arm (< −2). ‘Shallow amplifications’ and ‘shallow deletions’ were identified from samples with GISTIC indicators of +1 and −1 respectively.

### Defining Clock^Loss^ and Clock^Gain^ gene sets and calculating clock and hypoxia scores

Recurrently deleted/amplified genes were identified from genes that were deleted/amplified in at least 20% of samples within a cancer type and in at least one-third of cancers (> 7 cancers). Putative loss-of-function genes (Clock^Loss^) were defined as genes that were recurrently deleted and downregulated in tumour versus non-tumour samples. Putative gain-of-function genes (Clock^Gain^) were defined as genes that were recurrently amplified and upregulated in tumour versus non-tumour samples. Clock^Loss^ genes were CLOCK, CRY2, FBXL3, FBXW11, NR1D2, PER1, PER2, PER3, PRKAA2, RORA and RORB. Clock^Gain^ genes were ARNTL2 and NR1D1. For each patient, Clock^Loss^ and Clock^Gain^ scores were calculated from the mean log2 expression values of genes within each set. Molecular assessment of tumour hypoxia was performed using a 52-hypoxia gene signature(30). Hypoxia scores were determined from the mean log2 expression of the 52 genes.

### Survival, differential expression and multidimensional scaling analyses

We have published detailed methods for the aforementioned analyses(31), hence, the methods will not be repeated here. Briefly, for survival analyses, patients were separated into survival quartiles based on their Clock^Loss^ and Clock^Gain^ scores. Cox proportional hazards regression, Kaplan-Meier and receiver operating characteristic analyses were performed using the R survcomp, survival and survminer packages according to previous methods. For analyses in figure 5 investigating the crosstalk between hypoxia and the circadian clock, patients were separated into four groups based on their median clock and hypoxia scores. Nonparametric Spearman’s rank-order correlation analyses were performed to determine the relationship between clock and hypoxia scores in figure 5. Circular heatmaps in figure 6 were generated from Clock^Loss^ scores and HIF-target genes (CA9, VEGFA and LDHA) log2 expression values in glioma patients. Clock^Loss^ scores were ranked from high (purple) to low (yellow) in the heatmap. HIF-target genes were ranked by decreasing order of Clock^Loss^ scores. Spearman’s correlation analyses were performed on Clock^Loss^ scores and HIF-target genes expression values. Differential expression analyses between tumour and non-tumour samples and between the 4th and 1st quartile patients determined using Clock^Loss^ and Clock^Gain^ scores were performed using the Bayes method and linear model followed by Benjamini-Hochberg procedure for adjusting false discovery rates. Multidimensional scaling analyses in figure 2 were performed using the R vegan package (Euclidean distance) and permutational multivariate analysis of variance (PERMANOVA) was used to determine statistical difference between tumour and non-tumour samples.

**Fig. 1.**
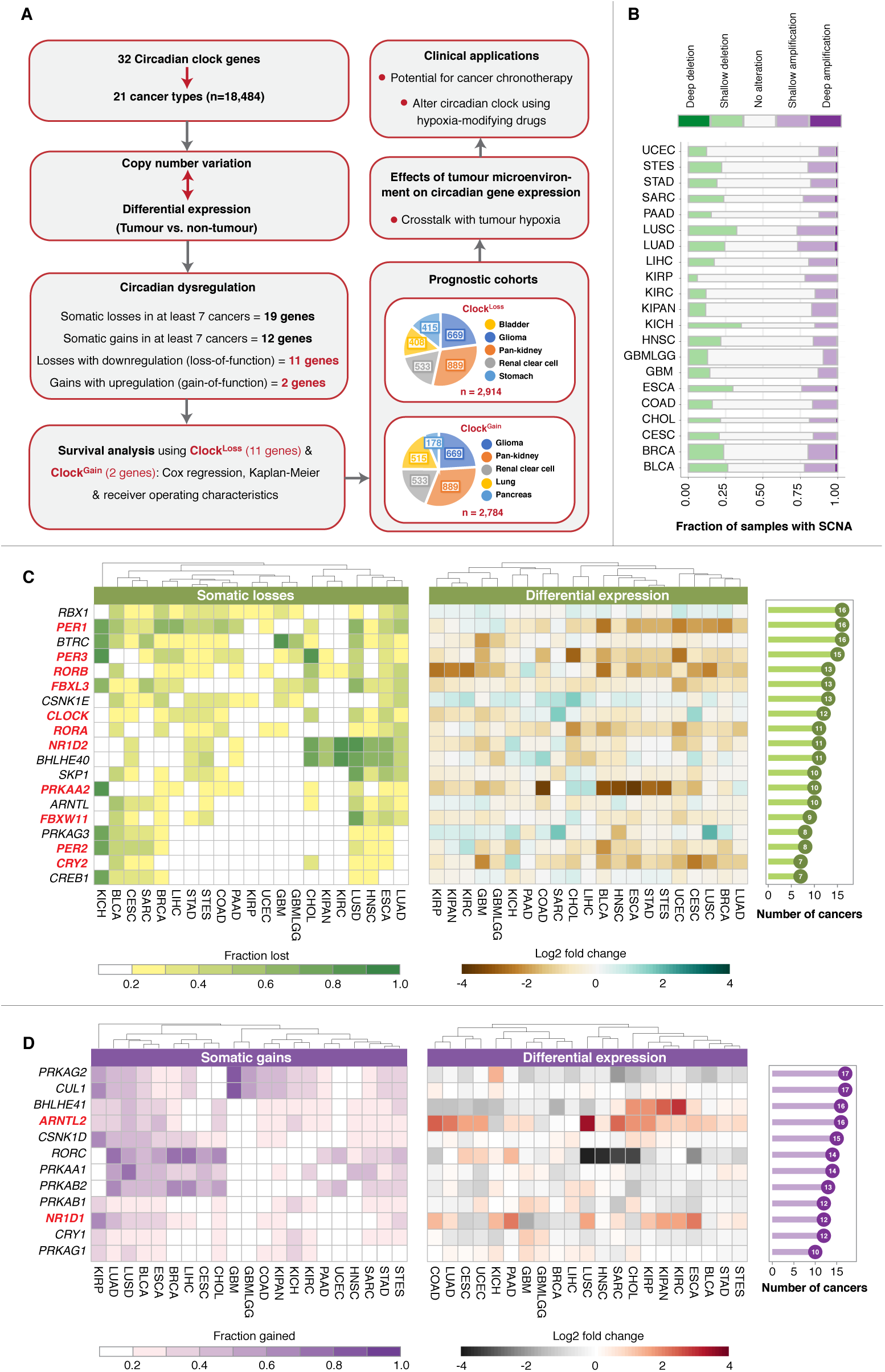
Circadian reprogramming in diverse cancer types. (A) Schematic diagram depicting the project design and the identification of putative loss-of-function and gain-offunction clock genes. Somatic copy number alteration (SCNA) and transcript expression of 32 clock genes are investigated in 21 cancer types. A total of 19 or 12 genes are recurrently lost or gained respectively. Of these SCNA events, 11 or 2 genes are also downregulated or upregulated in tumours, representing Clock^Loss^ and Clock^Gain^ gene sets respectively. Both gene sets are prognostic in seven cancer cohorts. Pie slices indicate the number of patients within each cancer type. Crosstalk between circadian genes and tumour hypoxia is investigated. (B) The proportion of samples with deep and shallow somatic alterations are represented using stacked bar graphs. The number of samples within each cancer type is represented by the width of the stacked bars. (C) Somatic losses and differential expression profiles of 19 clock genes that are recurrently deleted in at least 7 cancer types. (D) Somatic gains and differential expression profiles of 12 clock genes that are recurrently amplified in at least 7 cancer types. Bar charts on the far right represent the number of cancers with at least 20% of samples affected by copy number alteration. Heatmaps on the far left depict the cohort fraction in which a given gene is deleted or amplified. Cancer types are ordered using Euclidean distance metric. Heatmaps in the centre represent differential expression values between tumour and non-tumour samples. Clock^Loss^ and Clock^Gain^ genes are highlighted in red. Cancer abbreviations are listed in Additional file 2.

**Fig. 2.**
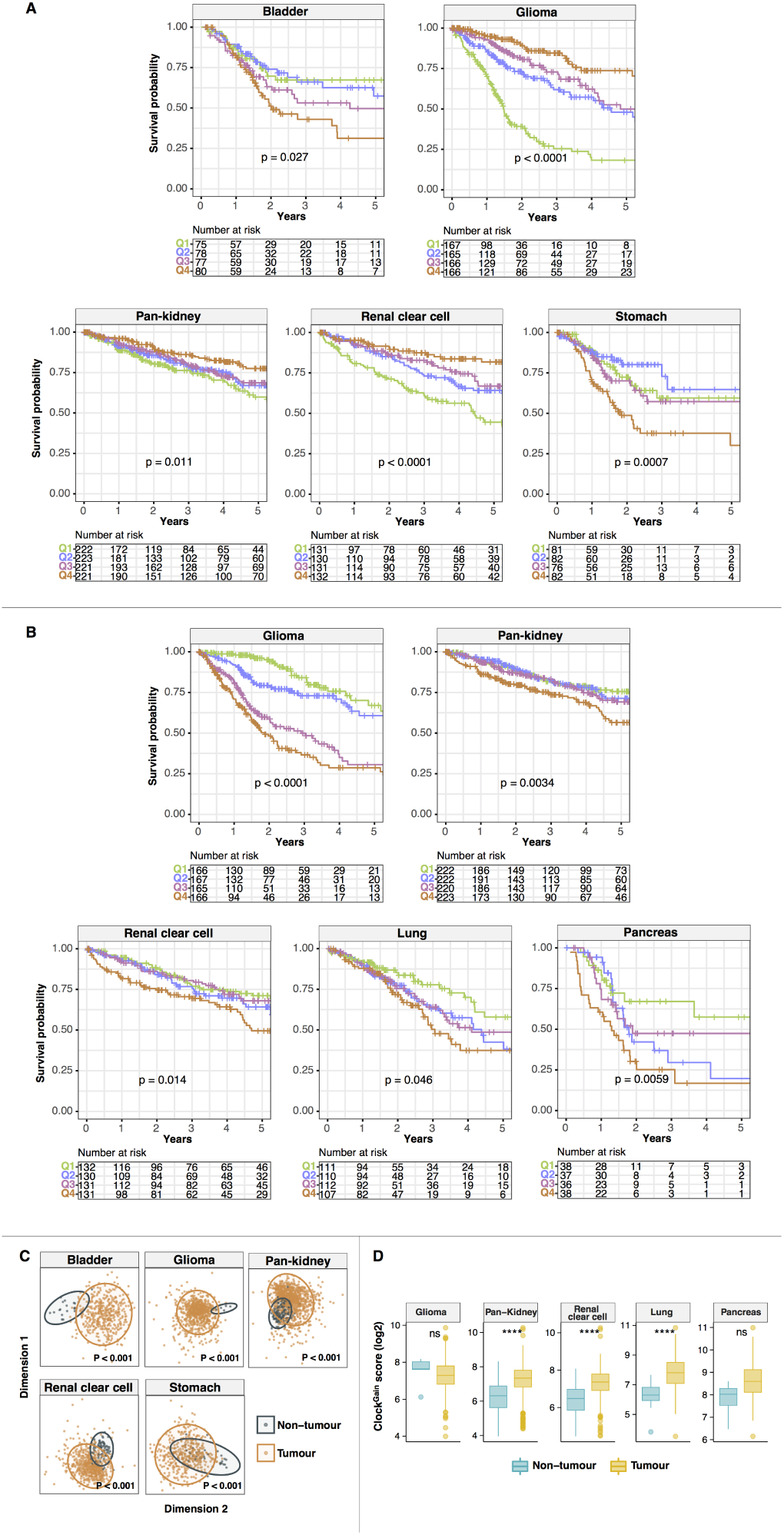
Prognostic significance of Clock^Loss^ and Clock^Gain^. Kaplan-Meier plots are generated using (A) Clock^Loss^ and (B) Clock^Gain^. Patients are quartile stratified based on their clock gene scores. P values are obtained from log-rank tests. (C) Ordination plots of multidimensional scaling analyses using Clock^Loss^ genes reveal significant differences between tumour and non-tumour samples. P values are obtained from PERMANOVA tests. (D) Expression distribution of Clock^Gain^ scores in tumour and non-tumour samples with statistical analyses performed using MannWhitney-Wilcoxon tests. P values are represented by **** < 0.00001. ns: nonsignificant.

### Pathway enrichment and transcription factor analyses

Differentially expressed genes (DEGs) were fed into GeneCodis(32) and Enrichr(33, 34). Using GeneCodis, DEGs were mapped against the KEGG and Gene Ontology databases. To determine whether DEGs were enriched for targets of stem cell-related transcription factors, Enrichr was used to map DEGs to ENCODE and ChEA chromatin immunoprecipitation sequencing profiles.

All plots were generated using R pheatmap and ggplot2 packages.

## Results

### Integrated genomic and transcriptomic analyses reveal conserved patterns of putative loss-and gain-of-function mutations in circadian clock genes

We retrieved 32 genes representing core circadian clock components from Kyoto Encyclopaedia of Genes and Genomes (KEGG) (Additional file 1). To determine the extent of circadian dysregulation across diverse malignancies, we analysed tumour copy number and mRNA differential expression profiles (tumour versus non-tumour) of 18,484 samples across 21 cancer types (Additional file 1) (Fig. 1A). Chromophobe renal cell carcinoma (KICH) exhibited the highest fraction of samples harbouring deleted clock genes (Fig. 1B). This was in contrast with another kidney cancer subtype, papillary renal cell carcinoma (KIRP), which had the lowest frequency of somatic deletions (Fig. 1B). When considering somatic amplifications, we observed that this was the highest in lung squamous cell carcinoma (LUSC) and the lowest in glioma (GBMLGG) (Fig. 1B). At an individual gene level, somatic deletions were observed in 19 circadian clock genes in at least 20% of samples within a cancer type and in at least one-third of cancer types (> 7 cancers) (Fig. 1A, 1C). Global somatic losses were observed in core circadian pacemaker genes, PER1, PER2, PER3, ARNTL (BMAL1), CLOCK, NR1D2 (REV-ERBB), RORA and RORB (Fig. 1C). For instance, PER1 was deleted in 16 cancers, while PER3 and CLOCK were lost in 15 and 12 cancers respectively (Fig. 1C). On the other hand, a distinct set of 12 genes exhibited global patterns of somatic gains, which included core clock genes namely ARNTL2 (BMAL2), CRY1, NR1D1 (REV-ERBA) and RORC (Fig. 1D). ARNTL2 was one of the most amplified genes found in 16 cancers, followed by RORC in 14 cancers and NR1D1 and CRY1 in 12 cancers each (Fig. 1D). Somatic copy number alterations (SCNAs) associated with differential transcript expression may represent putative loss- or gain-of-function events. Somatic losses accompanying transcript downregulation in tumours could indicate a loss-of-function and vice versa, somatic gains linked to transcript upregulation could imply a gain-of-function. Differential expression analyses between tumour and non-tumour samples were performed to identify genes that were significantly altered in tumours (> 1.5 fold-change, P<0.05) of at least 7 cancer types (Fig. 1C, 1D). Of genes that exhibited SCNAs, 11 and 2 genes were associated with putative loss-of-function and gain-of-function phenotypes respectively (Fig. 1A, 1C, 1D).

### Tumour suppressive and oncogenic potential of the circadian clock are context dependent

Given their global patterns spanning multiple cancer types, we reason that putative loss- and gain-of-function phenotypes would impact patient prognosis. We hypothesize that loss-of-function genes could have tumour suppressive qualities where gene deletions may give rise to cancer. On the contrary, gain-of-function genes are likely to harbour tumour promoting properties. If this is true, patients with low expression of loss-of-function genes (Clock^Loss^) would have poorer clinical outcomes. Likewise, high expression of gain-of-function genes (Clock^Gain^) would be associated with more advanced disease states and poorer outcomes. To evaluate the effects of Clock^Loss^ and Clock^Gain^ on overall survival, each patient was assigned a score based on the average expression values of 11 and 2 genes respectively: Clock^Loss^ genes (CLOCK, CRY2, FBXL3, FBXW11, NR1D2, PER1, PER2, PER3, PRKAA2, RORA and RORB); Clock^Gain^ genes (ARNTL2 and NR1D1). For Kaplan-Meier analyses, patients were separated into survival quartiles based on their Clock^Loss^ and Clock^Gain^ scores. Intriguingly, we found that both gene sets conferred prognostic information in seven cancer cohorts, allowing the stratification of patients into risk groups based on circadian dysregulation (Fig. 2A, 2B). Clock^Loss^ was prognostic five cohorts: bladder (P=0.027), glioma (P<0.0001), pan-kidney (consisting of chromophobe renal cell, clear cell renal cell and papillary renal cell carcinoma; P=0.011), clear cell renal cell (P<0.0001) and stomach (P=0.0007) (Fig. 2A). Likewise, Clock^Gain^ was also prognostic in glioma (P<0.0001), pan-kidney (P=0.0034) and clear cell renal cell (P=0.014) cohorts and two additional cohorts; lung (P=0.046) and pancreas (P=0.0059) (Fig. 2B).

Interestingly, we observed that patients within the 4th quartile (highest Clock^Loss^ scores) had the lowest mortality rates in glioma (hazard ratio [HR]=0.188, P<0.0001), pan-kidney (HR=0.520, P=0.001) and clear cell renal cell (HR=0.292, P<0.0001) cohorts (Table 1). This supports our initial hypothesis that downregulation/loss-of-function of putative tumour suppressive clock genes were linked to adverse clinical outcomes while patients with high expression of these genes would perform better. However, this was not the case for bladder (HR=2.081, P=0.0093) and stomach (HR=2.155, P=0.0054) cancers, where patients with high Clock^Loss^ scores (4th quartile) had increased death risks, suggesting that the function of Clock^Loss^ is tumour-type dependent (Table 1). In terms of Clock^Gain^, high expression levels were consistently associated with increased mortality rates in all five cohorts, supporting the hypothesis on tumour-promoting effects of these genes: glioma (HR=3.961, P<0.0001), pan-kidney (HR=1.890, P=0.00066), clear cell renal cell (HR=1.755, P=0.0062), lung (HR=2.023, P=0.006) and pancreas (HR=3.034, P=0.0022) (Table 1).

Given the prognostic significance of Clock^Loss^ and Clock^Gain^, we predict that their expression profiles would differ between tumour and non-tumour samples in these cancers. Indeed, as confirmed by multidimensional scaling analyses using Clock^Loss^, there was a clear separation between tumour and non-tumour samples in all five cohorts, suggesting that circadian dysregulation is a hallmark of cancerous cells (Fig.2C). Since Clock^Gain^ only involved two genes, we employed the Mann-Whitney-Wilcoxon test to compare the distribution of Clock^Gain^ scores in tumour and non-tumour samples. Clock^Gain^ was significantly upregulated in pan-kidney (P<0.00001), clear cell renal cell (P<0.00001) and lung cancer cohorts (P<0.00001) (Fig.2D). Glioma and pancreatic cancer cohorts had limited number of non-tumour samples, five and four samples respectively. Because of this, we did not observe any significant upregulation of Clock^Gain^ in these tumours (Fig.2D).

### Clock^Loss^ and Clock^Gain^ are independent prognostic factors

Multivariate Cox proportional hazards regression was used to determine whether Clock^Loss^ and Clock^Gain^ were independent of other clinicopathological features. Despite accounting for tumour, node and metastasis (TNM) staging, both gene sets remained independent predictors of overall survival; Clock^Loss^: bladder (HR=1.776, P=0.043), pan-kidney (HR=0.569, P=0.0055), clear cell renal cell (HR=0.433, P=0.00085), stomach (HR=2.070, P=0.0084) and Clock^Gain^: pan-kidney (HR=1.941, P=0.00054), clear cell renal cell (HR=1.856, P=0.0032), lung (HR=1.832, P=0.018) and pancreas (HR=2.890, P=0.0042) (Table 1). The glioma cohort consisted of low- and high-grade sub-types. Clock^Gain^ remained a prognostic factor in histological subtypes of astrocytoma (HR=3.048, P=0.0018) and oligodendroglioma (HR=2.764, P=0.018) (Table 1). Since both gene sets were independent of tumour stage, we evaluated their ability to improve the resolution of TNM staging. Kaplan-Meier analyses revealed that further delineation of risk groups within similarly staged tumours is afforded by both gene sets; Clock^Loss^: bladder (P<0.0001), pan-kidney (P<0.0001), clear cell renal cell (P<0.0001), stomach (P=0.024) (Fig. 3A) and Clock^Gain^: pan-kidney (P<0.0001), clear cell renal cell (P<0.0001), lung (P<0.0001), pancreas (P=0.048), astrocytoma (P<0.0001) and oligodendroglioma (P=0.023) (Fig. 3B).

**Fig. 3.**
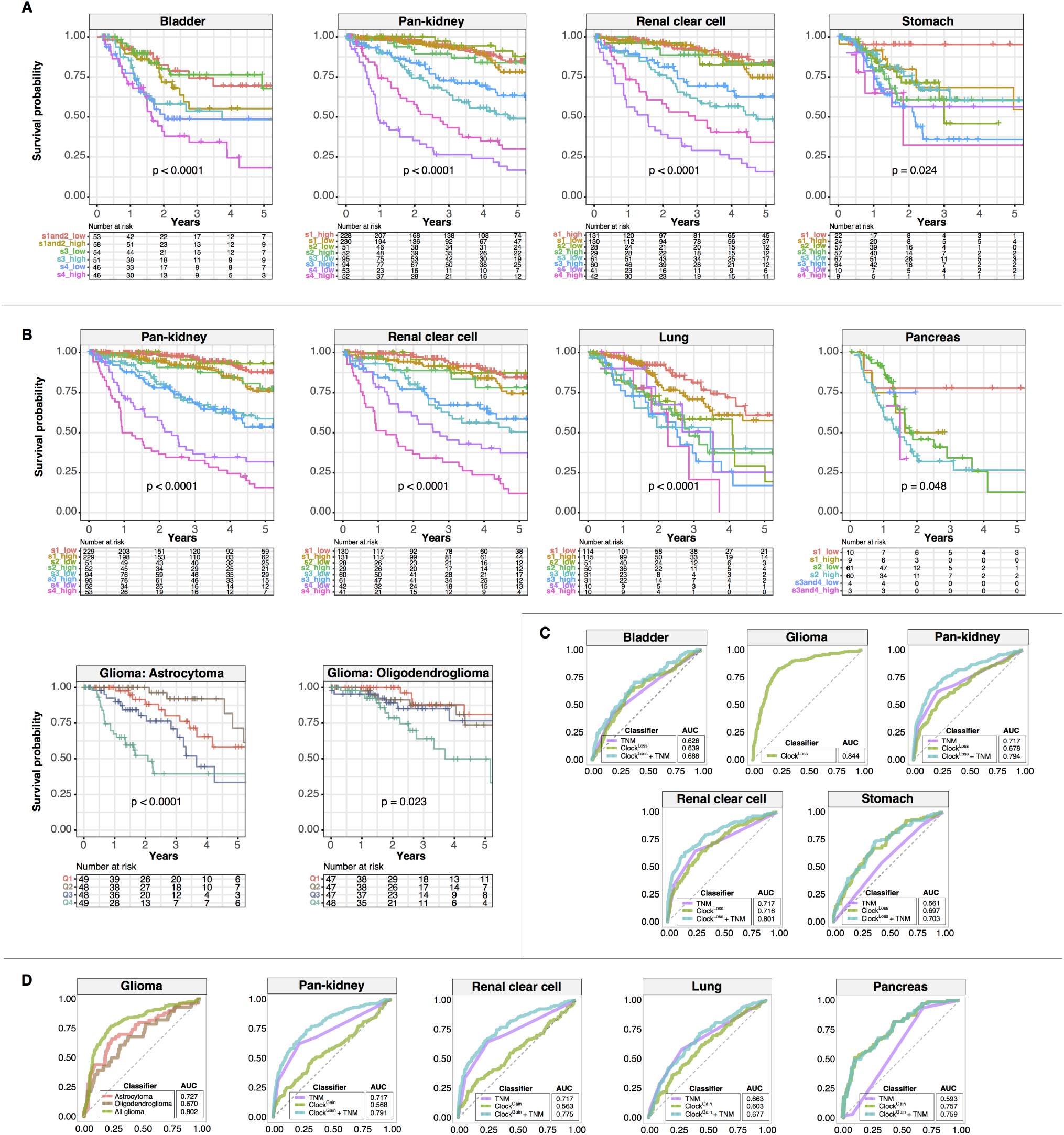
Clock^Loss^ and Clock^Gain^ are independent of tumour stage. Kaplan-Meier plots are generated from patients stratified according to TNM stage and (A) Clock^Loss^ and (B) Clock^Gain^. TNM staging is first used to stratify patients, followed by median stratification into highand low-score groups using Clock^Loss^ or Clock^Gain^. (B) Glioma histological subtypes, astrocytoma and oligodendroglioma, are quartile stratified using Clock^Gain^. P values are obtained from log-rank tests. ROC analyses on (C) Clock^Loss^ and (D) Clock^Gain^ to determine the specificity and sensitivity of both gene sets in predicting 5-year overall survival rates. ROC curves generated from clock gene sets are compared to those generated from TNM staging. AUCs for TNM stage are in accordance with previous work utilising TCGA datasets(35–37).

Receiver operating characteristic (ROC) analyses were employed to determine the predictive performance of Clock^Loss^ and Clock^Gain^ in comparison to TNM staging. Circadian gene sets were superior to TNM staging in predicting 5-year overall survival; Clock^Loss^: bladder (area under the curve [AUC]=0.639 vs. 0.626), stomach (AUC=0.697 vs. 0.561) and Clock^Gain^: pancreas (AUC=0.757 vs. 0.593) (Fig. 3C and 3D). Importantly, when clock gene sets and TNM staging were considered as a combined model, predictive performance was greater than clock genes or TNM when measured separately; Clock^Loss^: bladder (AUC=0.688), pan-kidney (AUC=0.794), clear cell renal cell (AUC=0.801), stomach (AUC=0.703) and Clock^Gain^: pan-kidney (AUC=0.791), clear cell renal cell (AUC=0.775), lung (AUC=0.677) and pancreas (AUC=0.759) (Fig. 3C and 3D). Within the glioma cohort, AUCs for Clock^Loss^ and Clock^Gain^ were 0.844 and 0.727 respectively (Fig. 3C and 3D). Clock^Gain^ was a prognostic indicator in glioma subtypes and ROC analyses confirmed its predictive performance in astrocytoma (AUC=0.727) and oligodendroglioma (AUC=0.670) (Fig. 3D).

### Dysregulated circadian timekeeping is associated with malignant progression

To further investigate the underpinning biological consequences of circadian clock dysregulation and determine how they link to unfavourable patient outcomes, we performed differential expression analyses on all transcripts to determine genes that were altered as a result of circadian perturbation by comparing patients from the 4th survival quartile to those from the 1st quartile. Interestingly, patients stratified using Clock^Loss^ had significantly higher number of differentially expressed genes (DEGs; −1.5 > log2 fold change > 1.5; P<0.01) (Fisher’s exact test, P= 2.2e-16) compared to Clock^Gain^ (Fig. 4A) (Additional file 3). Many DEGs were found to be in common between cancer types, more so for Clock^Loss^, suggesting the existence of conserved signalling cascades associated with circadian dysregulation in driving disease pathogenesis (Fig. 4A; Additional file 3). Gene Ontology (GO) and KEGG functional enrichment analyses revealed that circadian reprogramming of tumours resulted in the activation of a myriad of oncogenic pathways. Signalling pathways associated with cancer stem cell function (MAPK, Wnt, JAK-STAT, TGF-β and PPAR), metabolism, cell adhesion, cell proliferation, cell death, transmembrane transport and extracellular matrix organisation were among some of the most altered biological processes that likely underpin tumour aggression and decreased survival in these patients (Fig. 4B and 4C). Interestingly, with the exception of MAPK signalling, pathways associated with cancer stem cell function were only enriched in patients stratified using Clock^Loss^, suggesting that Clock^Loss^ genes are keenly linked to stem cell homeostasis (Fig. 4C). To further understand how DEGs were regulated, we analysed transcription factor (TF) binding using Enrichr and observed that DEGs were enriched for targets of TFs associated with self-renewal and cancer stem cell function (SUZ12, SOX2, REST, EZH2, SMAD4 and NANOG) (Fig. 4D). These TFs have previously been shown to promote metastasis, disease aggression and cancer stem cell maintenance(38–40).

**Fig. 4.**
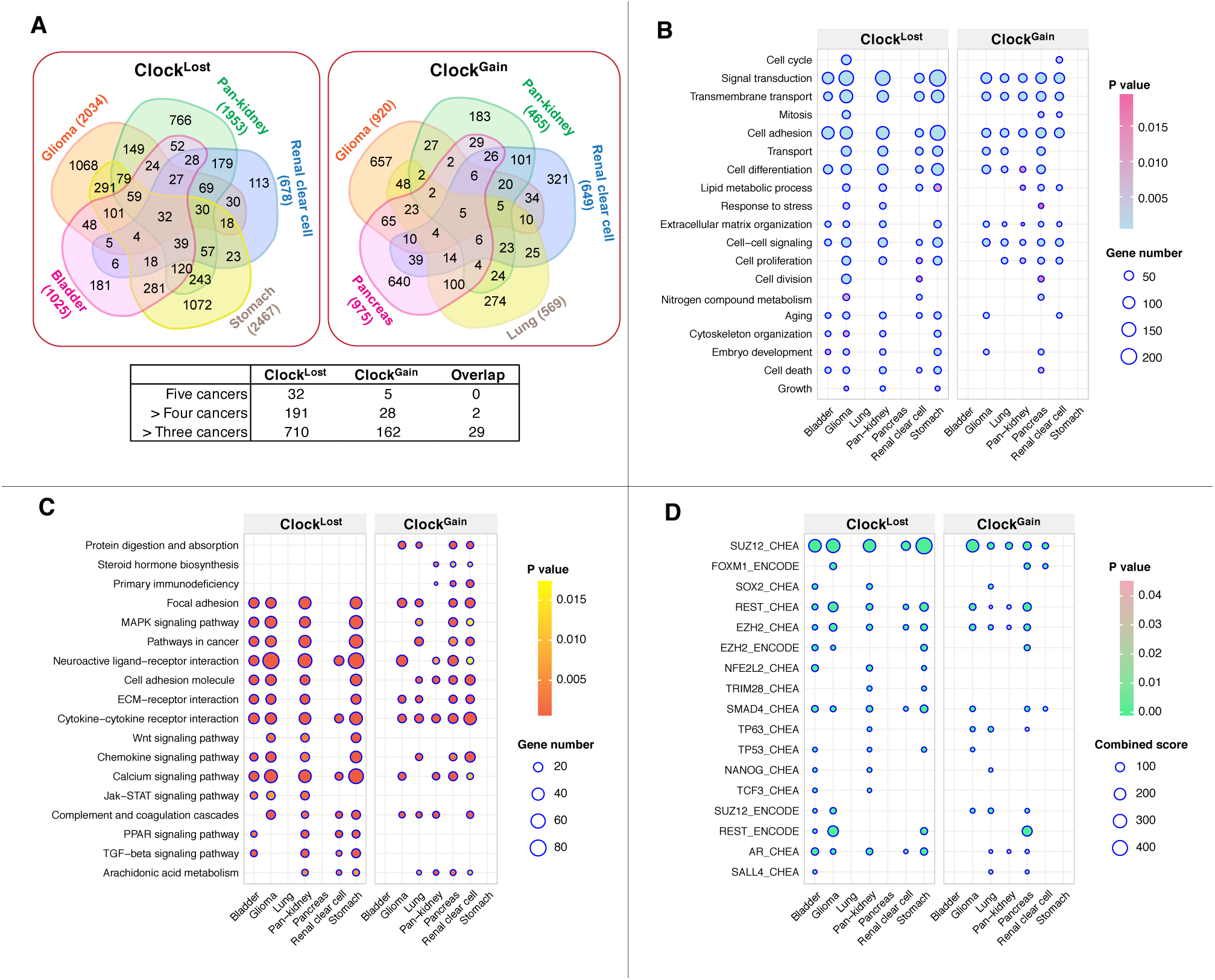
Circadian dysregulation drives malignant progression. Differential expression analyses are performed between 4th and 1st quartile patients determined using Clock^Loss^ or Clock^Gain^. (A) Venn diagrams illustrate the number of differentially expressed genes (DEGs) and their overlapping patterns in five cohorts. Numbers in parentheses represent DEGs. Table inset depicts the number of genes that are found to be in common in five, > four and > three cancer types. Overlap between common Clock^Loss^ and Clock^Gain^ genes are also depicted. Enriched (B) GO terms, (C) KEGG ontologies and (D) transcription factor binding associated with DEGs.

**Fig. 5.**
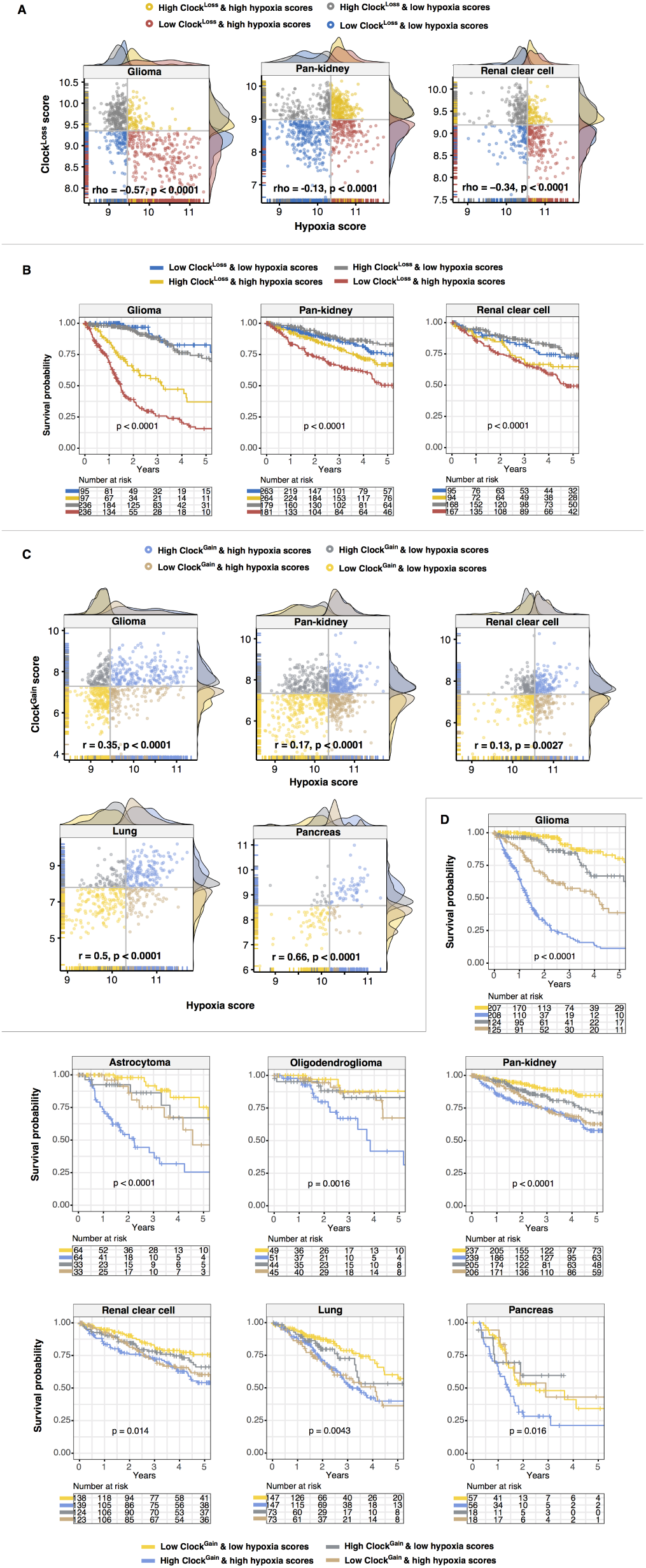
Prognostic relevance of the hypoxia and clock crosstalk. Scatter plots depict (A) significant negative correlations and (B) significant positive correlations between hypoxia and Clock^Loss^ or Clock^Gain^ scores respectively. Patients are grouped into four categories based on median clock and hypoxia scores. At the xand y-axes, density plots depict the distribution of clock and hypoxia scores. Kaplan-Meier analyses are performed on the four patient groups to determine the effects of crosstalk between hypoxia and (C) Clock^Loss^ and (D) Clock^Gain^ on overall survival in multiple cancers including glioma histological subtypes.

**Fig. 6.**
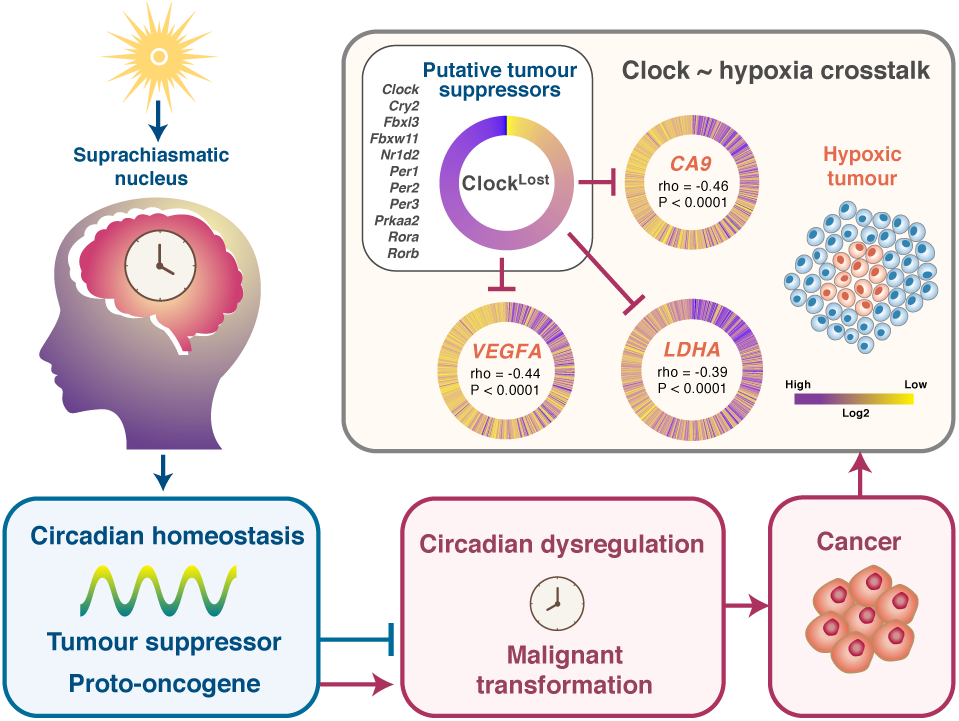
Model of the hypoxia-clock signalling axis in glioma. The circadian clock exerts tumour promoting or tumour suppressing qualities that are dependent on cellular types. Tumour suppressive Clock^Loss^ gene are negatively correlated with HIF-1A target genes (CA9, VEGFA and LDHA) in glioma. Clock^Loss^ scores are plotted such that each spoke of the circular heatmap represents individual patients that are sorted in descending order. Circular heatmaps for HIF-1A target genes are plotted with patients sorted in descending order of Clock^Loss^ scores. Spearman’s correlation coefficients between Clock^Loss^ and individual HIF-1A genes are depicted in the centre of the heatmap.

### Disease phenotypes of tumours with deranged circadian homeostasis are aggravated by hypoxia

Circadian oscillations of physiological processes such as metabolism, temperature and cortisol levels are affected by oxygen levels(41–43). Hypoxic responses are gated by the circadian clock and at the genomic level, BMAL1 and HIF-1A synergistically interact to co-regulate downstream genes(27). Crosstalk between hypoxia and the clock has profound implications on cancer pathophysiology(44, 45). We reason that tumour hypoxia could synergise with the circadian clock to impact disease progression. To determine the extent of the hypoxia-clock crosstalk, a hypoxia gene signature consisting of 52 genes was employed to calculate hypoxia scores in each patient(30). Remarkably, we observed significant negative correlations between hypoxia and Clock^Loss^ scores in glioma (rho=-0.57, P<0.0001), pan-kidney (rho=-0.13, P<0.0001) and clear cell renal cell (rho=-0.34, P<0.0001) cohorts (Fig. 5A). Patients were separated into four groups based on their hypoxia and Clock^Loss^ scores for survival analyses. Kaplan-Meier analyses employing the joint hypoxia-Clock^Loss^ model revealed that patients with more hypoxic tumours who concurrently had lower levels of Clock^Loss^ genes performed the worst: glioma (HR=7.218, P<0.0001), pan-kidney (HR=2.512, P<0.0001) and clear cell renal cell (HR=1.893, P=0.0054) (Fig. 5B) (Table 2). This observation is consistent with the predicted tumour suppressive roles of Clock^Loss^.

The trend is flipped when considering Clock^Gain^; significant positive correlations between hypoxia and Clock^Gain^ scores were observed in glioma (rho=0.35, P<0.0001), pan-kidney (rho=0.17, P<0.0001), clear cell renal cell (rho=0.13, P=0.0027), lung (rho=0.50, P<0.0001) and pancreas (rho=0.66, P<0.0001) (Fig. 5C). Consistently, patients with more hypoxic tumours and high Clock^Gain^ scores had the poorest survival outcomes: glioma (HR=9.210, P<0.0001), astrocytoma (HR=5.684, P<0.0001), oligodendroglioma (HR=4.085, P=0.0022), pan-kidney (HR=3.079, P<0.0001), clear cell renal cell (HR=1.877, P=0.0041), lung (HR=2.037, P=0.0012) and pancreas (HR=1.976, P=0.012) (Fig. 5D) (Table 2). Loss of tumour suppression or increase in tumour promoting properties resulted from circadian dysregulation is further exacerbated by hypoxia. The clock-hypoxia model may be used for delineation of patients into additional risk groups to support adjuvant therapy with hypoxia-reducing drugs in combination with mainstream chemotherapy and radiotherapy.

## Discussion

Disruption of circadian homeostasis is frequently observed in tumour cells. In a comprehensive study of circadian clock genes in 21 cancer types that takes into account genomic, transcriptomic and phenotypic (clinical prognosis) data, we demonstrated that clock genes were substantially altered by somatically acquired deletions and amplifications. Recurrent deletions or amplifications that were accompanied by altered transcript expression in tumours could represent novel loss-or gain-of-function phenotypes. To exploit these circadian targets in a clinical setting, we analysed survival outcomes using the Clock^Loss^ and Clock^Gain^ and confirmed the utility of both gene sets as prognostic tools in 2,914 and 2,784 patients involving seven diverse cancer cohorts.

Depending on cellular context, the circadian clock can exert both tumour-promoting or tumour-inhibiting properties. We observed that core clock genes, PERs, CRY2, CLOCK, NR1D2, RORA and RORB exhibited global patterns of somatic loss and downregulation across multiple tumour types (Fig. 1C). We demonstrated that loss-of-function of these genes resulted in increased death risks in patients, which highlight their protective roles (Fig. 2, 3). However, tumour suppressive qualities appear to be cancer type-specific; Clock^Loss^ genes were associated with adverse survival outcomes in bladder and stomach cancers (Fig. 2, 3). A study on breast cancer revealed that CpG methylation on PER promoters is responsible for PER deregulation in tumours(46). PER2 enhances estrogen receptor-α (ERα) degradation leading to growth inhibition of estrogen receptor-positive breast cancer(47). Studies on glioma(48), head and neck squamous cell carcinoma(49), lung(50), colorectal(51) and liver cancers(52) demonstrated that PERs CRYs and BMAL1 are downregulated in tumours and are likely to harbour tumour suppressing qualities. BMAL1 overexpression increases the sensitivity of colorectal cancer cells to oxaliplatin by ATM pathway activation(17). Overexpression of PER2 in pancreatic cancer cells inhibits cellular proliferation, increases apoptotic rates and had a synergistic effect with cisplatin(20). BMAL1 binds to TP53 promoter to induce cell cycle arrest and apoptosis in pancreatic cancer cells in a TP53-dependent fashion(53). Additional examples on tumour suppressive functions of clock genes are elegantly reviewed by Fu et al.(54).

We demonstrated that a non-overlapping subset of circadian clock genes, known as Clock^Gain^, were somatically amplified and upregulated in tumours. To our knowledge, most reports on circadian dysregulation in cancer have focused on the pacemaker’s tumour suppressive roles. Nonetheless, evidence on its oncogenic potential have started to emerge. CLOCK expression levels are higher in more aggressive ERα-positive compared to ERα-negative breast tumours and estrogen promotes the binding of ERα to estrogen-response elements in the CLOCK promoter(55). CRY2 is overexpressed in colorectal cancer samples that are resistant to chemotherapy(21). CRY2 is linked to poor survival outcomes and its silencing could increase sensitivity to oxaliplatin(21). Upregulation of PER2 and CRY1 in gastric cancer and CRY1 in colorectal cancer correlate with more advanced disease states and lymph node metastasis(56, 57). CLOCK expression is increased in high-grade glioma tissues and is required for glioma progression through modulation of NF-κB activity(58). CLOCK and BMAL1 are required for acute myeloid leukaemia cell growth and leukaemia stem cell maintenance(59).

Circadian reprogramming of tumours is closely associated with metabolic perturbations, activation of cell proliferation, induction of cancer stem cell self-renewal pathways (Wnt/βcatenin, JAK-STAT and TGF-β) and enrichment of binding targets of self-renewal TFs (Fig. 4). Circadian disruption in mouse xenograft models results in tumour progression through Wnt10A-dependent activation of angioand stromagenesis(60). BMAL1 overexpression promotes mouse embryonic fibroblast cell proliferation by Wnt signalling activation(61). In acute myeloid leukaemia, maintenance of circadian homeostasis is required for leukaemia stem cell self-renewal and inhibition of BMAL1 results in the downregulation of β-catenin and other TFs involved in self-renewal(59). BMAL1 loss also causes stem cell arrhythmia in squamous cell tumours(62). Taken together, the role of the circadian clock in stem cell homeostasis is likely to be conserved across multiple tissue types.

Equally important, we demonstrated that tumour hypoxia further aggravates the extent of circadian dysregulation resulting in increased death risks, suggesting that interactions between PER-ARNT-SIM components of both circadian and hypoxia pathways could synergistically influence disease progression. The expression of clock genes with putative tumour suppressive properties (Clock^Loss^) is negatively correlated with tumour hypoxia (Fig. 5A and 5B; Fig. 6). On the other hand, tumour promoting Clock^Gain^ genes were positively correlated with hypoxia (Fig. 5C and 5D). PER1 and CLOCK levels are elevated when mouse brain is exposed to hypoxia(63). VHL, a protein involved in proteasomal degradation of HIFs, is frequently mutated in renal cancers and consequently, this results in HIF accumulation leading to the induction of proangiogenic factors and malignant progression(64). HIF-1A promotes the amplitude of PER2 rhythms in renal cancer(65). Considering the high degree of sequence similarities between HIF-1A ([A/G]CGTG) and BMAL1 (CACGTG) binding motifs, it is not surprising that HIF-1A and BMAL1 co-occupy 30% of all genomic loci(27). Indeed, we observed significant negative correlations between Clock^Loss^ and HIF1A targets (CA9, VEGFA and LDHA) in glioma (Fig. 6). Reduction in tumour protective effects of Clock^Loss^ coupled with elevated HIF signalling could, together, help explain the significant increase in mortality rates in glioma patients (Fig. 6). These results are supported by another study on breast cancer where hypoxia is negatively correlated with PER2 expression and promotes PER2 degradation to stimulate epithelial-mesenchymal transition(66).

## Conclusion

One of the key strengths of the comparative approach we took was that it reveals non-mutually exclusive oncogenic and tumour suppressive properties of the circadian clock. Genes that confer tumour attenuating effects in one cancer type could very well play an opposing role in another cancer type. We confirmed prognostic values of two circadian gene sets in seven cancer cohorts including difficult-to-treat cancers such as glioma and pancreatic cancer; these genes may be prioritised as new therapeutic targets. Although prospective analysis is needed, we anticipate that our findings will provide an important framework for cancer chronotherapy initiatives(67–69) by enabling patient stratification based on circadian biomarkers to enhance therapeutic success. Moreover, therapeutic modification of the clock should help lessen damage caused by tumour hypoxia. Likewise, it will be important to investigate whether manipulating hypoxia levels could improve adjuvant chronotherapy when used in combination with first-line treatments.

